# Genotype-phenotype map of an RNA-ligand complex

**DOI:** 10.1101/2020.12.17.423258

**Authors:** Olga Puchta, Grzegorz Sobczyk, Vanessa Smer-Barreto, Hollie Ireland, Marc Vendrell, Diego A. Oyarzún, Janusz M. Bujnicki, Graeme Whyte, Grzegorz Kudla

## Abstract

RNA-ligand interactions play important roles in biology and biotechnology, but they often involve complex three-dimensional folding of RNA and are difficult to predict. To systematically explore the phenotypic landscape of an RNA-ligand complex, we used microarrays to investigate all possible single and double mutants of the 49-nt RNA aptamer Broccoli bound to the fluorophore DFHBI-1T. We collected more than seven million fluorescence measurements in varying conditions, and inferred dissociation rate constants, spectral shifts, and intragenic epistasis. Our results reveal an unexpectedly complex phenotypic landscape, in which mutations near the fluorophore binding pocket modulated magnesium-, potassium- and fluorophore-binding and fluorescence spectra, while distal mutations influenced structural stability and fluorescence intensity. We trained a machine learning model that accurately predicted RNA secondary structure from local epistatic interactions, despite the presence of G-quadruplexes and other noncanonical structures. Our experimental platform will facilitate the discovery and analysis of new RNA-ligand interactions.

## Introduction

In addition to its role as messenger in protein synthesis, RNA performs functions that depend on its three-dimensional folding and binding to other molecules. Such interactions are important in biology (eg, the specific recognition of tRNA sequence and structure by aminoacyl tRNA synthetases (Ibba and Soll, 2000) and biotechnology (eg, the inhibition of SARS-CoV ribosomal frameshifting by a ligand of the FSE pseudoknot (Park et al., 2011; Warner et al., 2018)). However, RNA-ligand interactions are difficult to measure, and computational methods are poor at predicting noncanonical RNA structures, such as G-quadruplexes, that often establish RNA-ligand specificity.

The key to accurate prediction of RNA structures and interactions is experimental data that connects RNA sequence to structural properties. Such data has been instrumental in developing commonly used secondary structure predictors, such as UNAfold and RNAfold (Lorenz et al., 2011; Markham and Zuker, 2008), and 3D structure predictors (Boniecki et al., 2016; Das and Baker, 2007). Unfortunately, the data used to train existing predictors has low coverage of noncanonical structures or RNA-ligand complexes, and little information about the effects of external conditions on structural stability. To address these gaps, here we performed a massively parallel assay of the effects of RNA sequence variation and external conditions on the binding of the 49-nt RNA aptamer Broccoli to the fluorophore DFHBI-1T (Filonov et al., 2014).

### Single mutation effects

We designed an oligonucleotide library that encodes wild-type Broccoli and all its 10,731 (3×49 + (3×49) × (3×48) / 2) single and double mutants. Each variant was attached to a stabilizing scaffold (Filonov et al., 2015) and to three or more unique 60-nt probes, allowing specific hybridisation to spots on an Agilent cDNA microarray (Fig. 1A, B). We transcribed the library with T7 polymerase and labeled a small fraction of RNA with Cyanine-3 (Cy3) as loading control. To evaluate the effects of mutations on fluorescence, we measured the green signal (aptamer-DFHBI-1T complex) and red signal (Cy3-labeled aptamer) using an automated fluorescence microscope. To guide the analysis of structure-function relationships, we predicted the 3D structure of Broccoli by homology modeling using the known crystal structure of Spinach (Warner et al., 2014).

**Figure 1.**
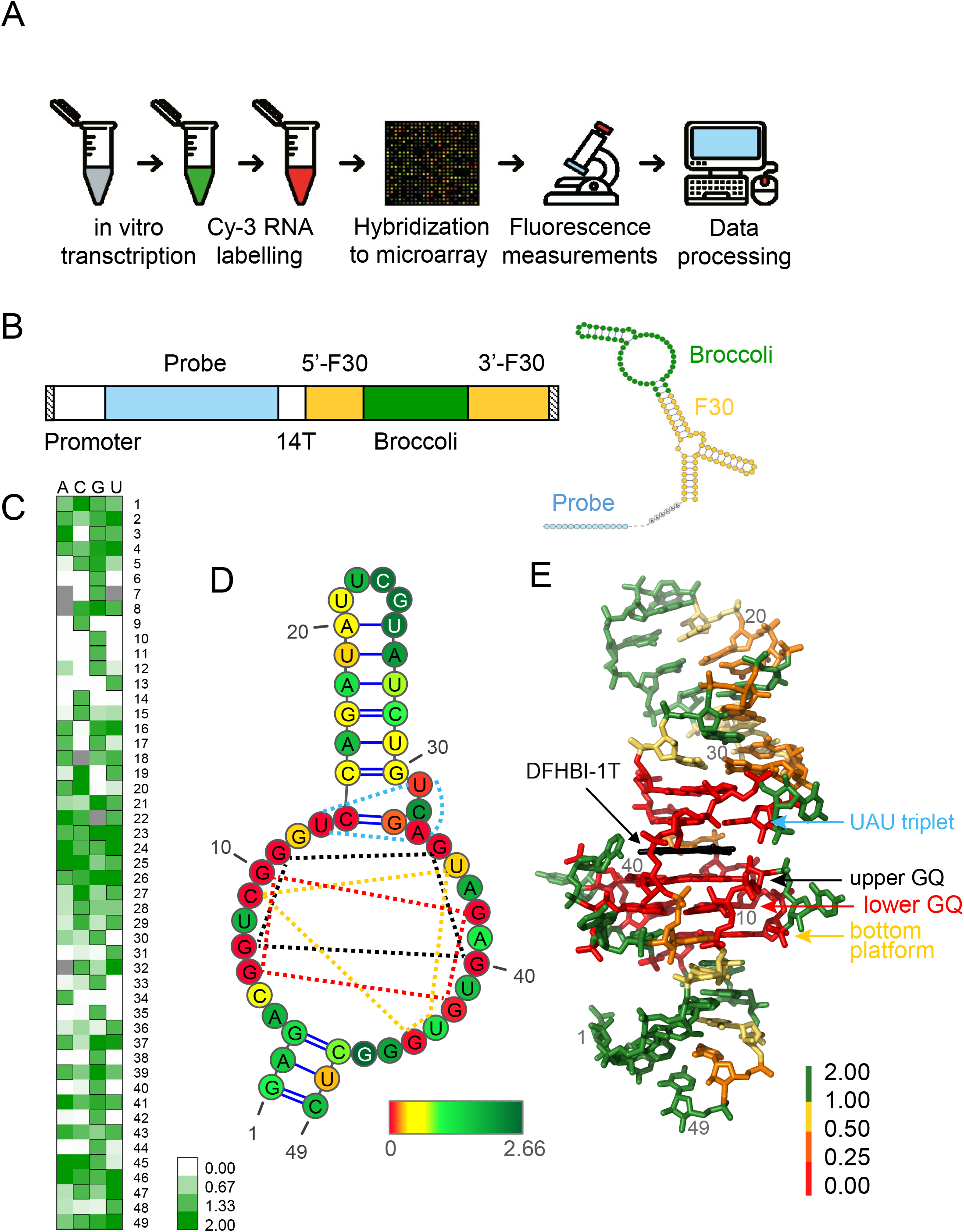
Genotype-phenotype mapping of Broccoli RNA. (A) Experimental design. (B) Design of oligonucleotide library. The bar on the left represents the DNA oligonucleotide library, the folded structure is the transcribed RNA. 5’F30 and 3’F30 indicate the 5’ and 3’ sides of the F30 scaffold, respectively. (C) Fluorescence of single mutants of Broccoli in 10 mM Mg^2+^, 140 mM K^+^, pH=5.5. Rows indicate positions along the RNA and columns indicate substitutions to one of the four bases: A, C, G or U. Black outlines indicate wild-type positions. Fluorescence of wild-type Broccoli is set as 1. (D-E) Median fluorescence of single mutants in each position mapped onto the secondary (D) and tertiary (E) structure models of Broccoli. The dashed lines in (D) indicate the U13-U31-A34 base triplet, and the G6-G10-G38-G42, G7-G11-G35-G40, and C9-U36-U43-G44 quadruples.

The microarray measurements were consistent between experimental replicates and with published data (Fig. S1). In most positions, mutations had mild-to-moderate negative effects on fluorescence, but some variants were brighter than the wild-type (Fig. 1). Although the sequence of the apical loop, UUCG, is known as a stabilizing tetraloop (Heus and Pardi, 1991), most mutations in this region increased fluorescence, as seen before (Ageely et al., 2016). Mutations in canonically (Watson-Crick C-G, A-U and G-U) paired regions typically reduced fluorescence, consistent with their predicted destabilizing effect. Unlike in previous reports (Ageely et al., 2016) mutations in the terminal stem were tolerated, presumably because our design comprised a stabilizing scaffold. Nucleotide G12, thought to be essential because it forms hydrogen bonds with the fluorophore (Warner et al., 2014), tolerated mutation to A, but not to C or U. Mutations in the two G-quadruplexes (GQ) that form the fluorophore binding platform, and in the U13-A33-U31 Hoogsteen base triple which seals the binding pocket from the top, reduced fluorescence by more than 90%, confirming the importance of these structural elements for fluorophore binding.

### Epistasis

Previous large-scale studies showed that the effects of mutations in a gene usually depend on the presence of other mutations in the same gene, a phenomenon known as intragenic epistasis (Domingo et al., 2018; Li et al., 2016; Puchta et al., 2016; Sarkisyan et al., 2016). To visualise epistasis, we compared the effects of single mutations in the wild-type background, and in each of the 147 (3 x 49) single-mutant backgrounds. Although mutational profiles showed similarities across multiple backgrounds, there were also clear differences, indicating the existence of intragenic epistatic effects (Fig. S2).

Epistasis can be partitioned into nonspecific (global) epistasis, where the phenotypes of double mutants can be predicted from the effect sizes of single mutations, and specific (local) epistasis, where phenotypes of double mutants also depend on the precise identity of both mutations (Domingo et al., 2019; Kondrashov and Kondrashov, 2001; Otwinowski et al., 2018). To calculate nonspecific epistasis, we fitted a locally weighted polynomial regression (LOESS) model to estimate the typical fluorescence of double mutants given the observed fluorescence of single mutants (Fig. 2A). For each double mutant, we then defined specific epistasis as the deviation between observed fluorescence and fluorescence predicted by the nonspecific epistasis model (Fig. S3, top right). Nonspecific epistasis explained 63% of the variation in fluorescence, compared to 42% of variation explained by an additive model (Fig. S4). Unlike the additive model, the nonspecific epistasis model correctly predicted non-negative fluorescence and lack of positive epistasis between pairs of large-effect mutations.

**Figure 2.**
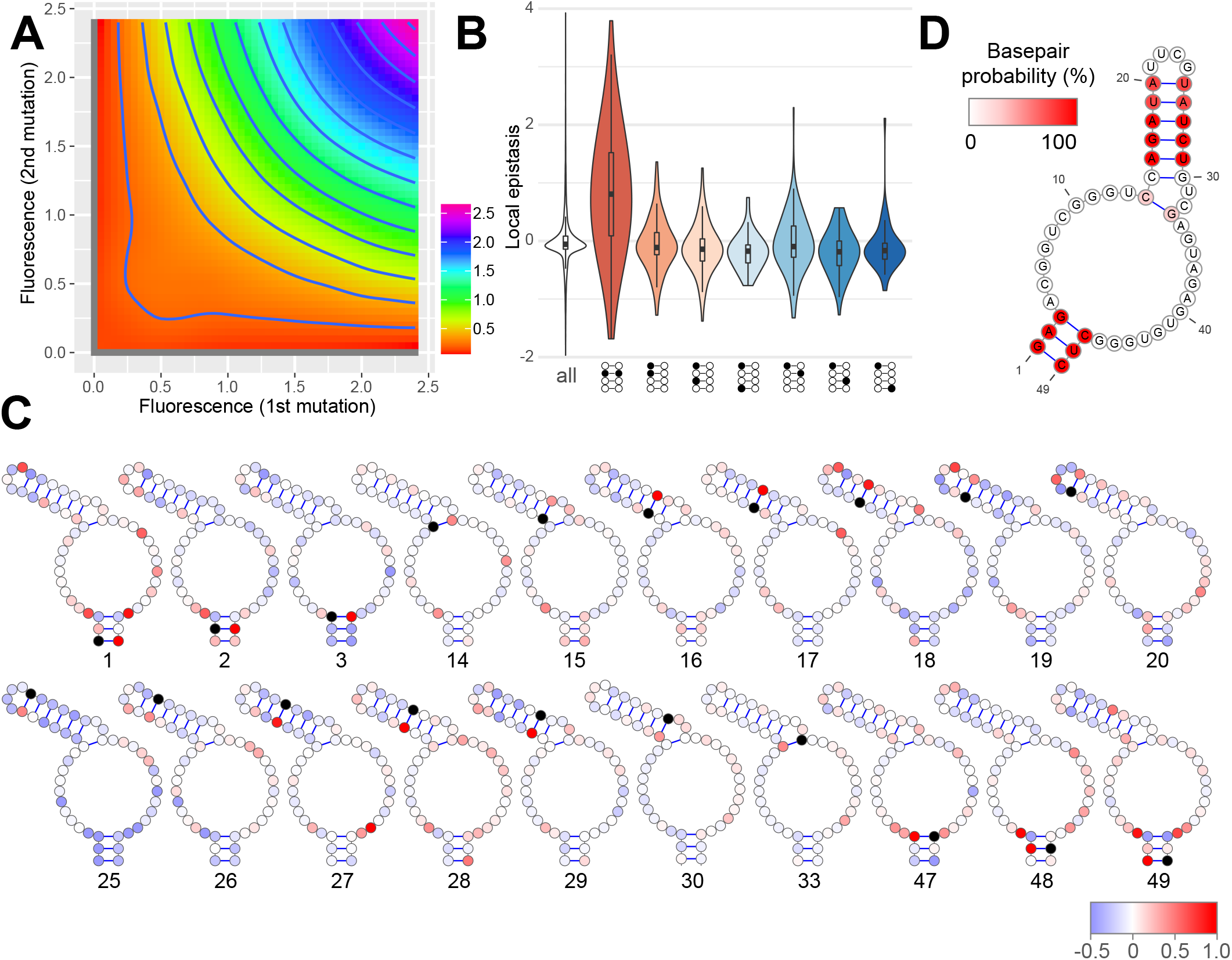
Local epistatic interactions predict RNA secondary structure. (A) Model of global epistasis. The colour represents the fluorescence of double mutants as a function of fluorescence of corresponding single mutants (on X and Y axes). The surface has been smoothed by locally weighted polynomial regression (LOESS). Experimental conditions: 10 mM Mg^2+^, 10 uM DFHBI-1T, 140 mM K^+^, pH=5.5, T=23°C. (B) Distribution of local epistasis between pairs of mutations found within secondary structure elements. Diagrams below the X axis show the mutual positions of the focal pair of mutations (filled circles) within a double-stranded region (empty circles). (C) Profiles of local epistasis of nucleotides involved in Watson-Crick basepairs. The position of the focal nucleotide is indicated by a black circle and a number below each structure. Red colour indicates positive epistasis and blue, negative epistasis. (D) Frequency for each base pair to be correctly identified as paired by the ensemble of SVM classifiers.

Local epistatic interactions measured by deep mutational scanning have been used to predict structures of proteins and RNAs (Rollins et al., 2019; Schmiedel and Lehner, 2019; Zhang et al., 2020). To evaluate the relation with RNA structure, we analysed local epistasis between pairs of nucleotides, depending on their position within the secondary and tertiary structures of Broccoli. Base-paired nucleotides showed strong positive epistasis, consistent with the restoration of function by compensatory mutations, whereas non-paired nucleotides located within the same stem showed negative epistasis (Fig. 2B,C, Fig. S3). Pairs in which one or both partners were part of a triplet or quadruplet structure typically showed weak epistasis, whereas loop nucleotides showed both positive and negative epistasis (Fig. S5A). Unexpectedly, proximity in the 3D structure was associated with lower absolute values of epistasis (Fig. S5B,C).

The association between epistasis and structural elements suggests that it may be possible to predict RNA structure from epistasis signals. To test this principle, we employed supervised machine learning to predict base pairings of the wild-type Broccoli. We trained support vector machine (SVM) classifiers on three features for each candidate base pair. The chosen features were drawn from local epistasis signals associated with each candidate pair (details in Methods) on the basis of their ability to discriminate between paired and non-paired candidates in the feature space (Fig S6). We labelled all pairs (N=990) as paired or non-paired according to the known wild-type structure. We employed stratified sampling on a 70-30 split between training and test data, to account for the class imbalance between paired and non-paired candidates (10:980 ratio). We trained an ensemble of 1000 classifiers that achieved average sensitivity of 76% and specificity of 98%.

Most pairs were accurately predicted by the SVM, with the exception of pair 15:30, which possibly indicates constraints on nucleotide identity (Fig. 2D). The minimum folding energy (MFE) structure comprises five base pairs not found in our 3D structural model, and none of these extra pairs were recovered by the SVM, consistent with the expectation that these pairs are not compatible with fluorescence (Fig. S7). Some candidate pairs (eg 4:46 and 5:45) were not present in the MFE structure, but were consistently called as paired by the SVM (Fig. S3). These base pairs, which have been reported as paired in some previous studies (Ageely et al., 2016), may represent alternative functional conformations of Broccoli. Consistently, pairs 4:46 and 5:45 also showed some propensity for base-pairing in 3D folding simulations.

### Gene-environment interactions

Our approach allows the estimation of multiple molecular phenotypes through deep mutational scanning across experimental conditions. We performed 168 experiments with varying fluorophore concentrations, Mg^2+^ and K^+^ ion concentrations, pH, temperature, and excitation/emission wavelengths. While the red signal, corresponding to the amount of RNA hybridized to each spot, was similar across experiments, the intensity of green signal depended on the experiment. Assays performed in similar conditions yielded highly correlated fluorescence profiles (Fig. S8). As expected, high concentrations of fluorophore, Mg^2+^ or K^+^ in the buffer were associated with increased brightness of most variants (Fig. 3, S9).

**Figure 3.**
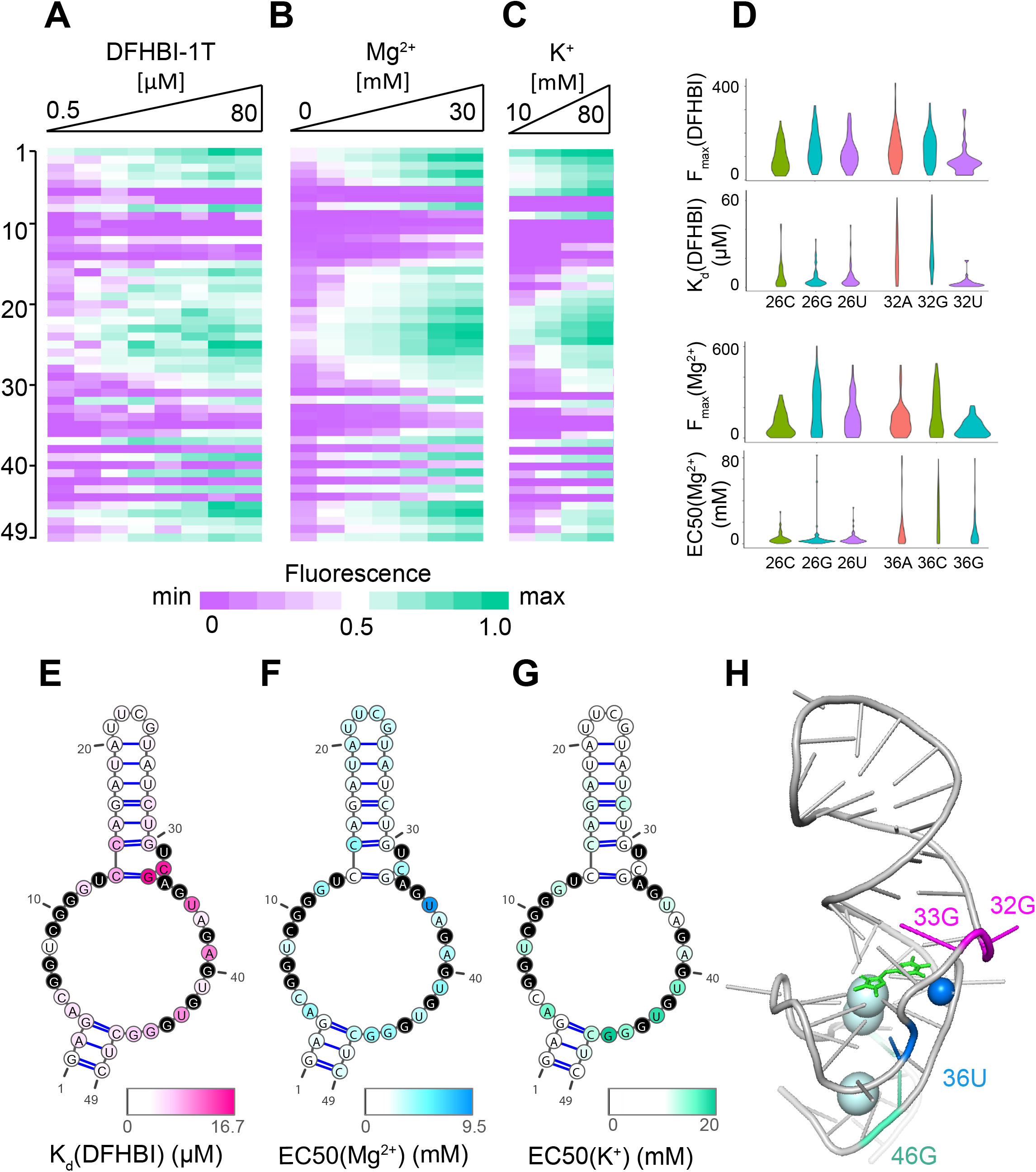
Dependence of mutational effects on environmental conditions. (A-C) Fluorescence of Broccoli mutants as a function of mutation (Y axis) and concentration of DFHBI-1T (A), Mg^2+^ (B), or K^+^ (C). Each data point represents the median fluorescence of single mutants (in the Mg^2+^ gradient), or the median fluorescence of single mutants and a subset of double mutants (in the DFHBI-1T and K^+^ gradients) (see Methods). (D) Distributions of K_d_(DFHBI-1T), F_max_(DFHBI-1T), EC50(Mg^2+^), and F_max_(Mg^2+^) of all variants with a mutation at the indicated position, in single mutants and a subset of double mutants (see Methods). (E-G) Median K_d_(DFHBI-1T) (E), EC50(Mg^2+^) (F), and EC50(K^+^) (G) of single mutants mapped onto the secondary structure. (H) Positions of mutations that cause the largest increase of K_d_(DFHBI-1T) (pink) EC50(Mg^2+^) (blue), and EC50(K^+^) (cyan). DFHBI-1T is shown in green, a Mg^2+^ ion is in dark blue, and K^+^ ions are in cyan.

Fluorescence measurements in varying fluorophore concentrations provide information about the strength of RNA-fluorophore associations across the library. We calculated the DFHBI-1T dissociation constant (Kd) and maximum fluorescence (F_max) for variants of Broccoli by fitting a Hill equation: F(x)= F_max*[L]/([L]+Kd), with Hill coefficient equal to 1 to represent a single binding site. Mutations with the largest effect on fluorophore affinity were found in the fluorophore binding pocket, in positions C32 and G33 (Fig. 3). Residue A69 in Spinach influences fluorophore access into its binding site (Warner et al., 2014); consistently, mutations of the homologous residue A39 in Broccoli decreased affinity. F_max correlated well with fluorescence measured in individual experiments, but weakly with Kd (Fig. S10). Some mutations (such as C32U) increased DFHBI-1T-Broccoli affinity and decreased fluorescence, while others (eg. C32A and C32G) had an opposite effect (Fig. 3D). This counterintuitive observation may be explained by postulating that weak binding reduces photobleaching and increases the proportion of RNA-fluorophore complexes found in an active state. Indeed, a previous study described fast fluorescence decay and fluorophore-dependent recovery of RNA aptamers, which has been attributed to accelerated photoconversion of the fluorophore when bound to the RNA (Han et al. 2013). Wild-type Broccoli had one of the highest affinities to the fluorophore, but many mutants were brighter than wild-type (Fig. S10, see also (Ketterer et al., 2015), consistent with the original selection strategy which relied primarily on DFHBI binding rather than fluorescence (Filonov et al., 2014).

Similar to DFHBI-1T titration, Mg^2+^ and K^+^ titration showed monotonically increasing relationships between ligand concentration and fluorescence (Fig. 3, S9, S10, S11), and we used the Hill equation to calculate EC50(Mg^2+^) and EC50(K^+^), the ion concentrations at which half of maximum fluorescence was observed. Broccoli was initially selected in low-magnesium conditions to improve *in vivo* fluorescence (Filonov et al., 2014), and wild-type Broccoli showed low magnesium sensitivity. On average, mutants required ~1.8 times as much magnesium as the wild-type to achieve half-maximum fluorescence, but variants mutated in position 36 required ~5.3 times as much magnesium (Fig. 3C, D). In the structural model, nucleotide U36 is close to a coordinated Mg^2+^ ion, suggesting a role in binding (Fig. 3E). Mutations in positions 43 and 46 increased sensitivity to K^+^ concentration, suggesting a role in the formation of the G-quadruplex structures, which are stabilized by K^+^ ions. While changes of magnesium and potassium concentration affected the fluorescence signal differently in different mutants, changes of temperature and pH showed less interaction with individual mutations (Fig. S9).

### Fluorescence spectra

Fluorescent RNA aptamers are known to change emission and excitation spectra when bound to different fluorophores (Chen et al., 2019; Song et al., 2014; Steinmetzger et al., 2019), or when mutated in specific positions (Filonov et al., 2019; Warner et al., 2014). To check the influence of mutations on spectral properties of Broccoli, we imaged the library using GFP (green) and CFP (cyan) filter sets. Green and cyan fluorescence were strongly correlated for almost all variants, except two groups of outliers (Fig. 4A). A reduced cyan signal was found exclusively in variants that carried the G12A substitution (Fig. 4B,C), while enhanced cyan fluorescence was associated with mutations that disrupted Watson-Crick base-paring in the fluorophore-proximal side of the upper stem (Fig. 4 B,C). As reported previously (Warner et al., 2014), mutations in the U-A-U base triple atop the fluorophore greatly reduced fluorescence and caused a blue-shift of the spectrum. Position A39, previously reported as a spectral tuning spot for DFHO-Broccoli_29-1 red/orange complex (Filonov et al., 2019), had no effect on the fluorescence spectrum of DFHBI-1T.

**Figure 4.**
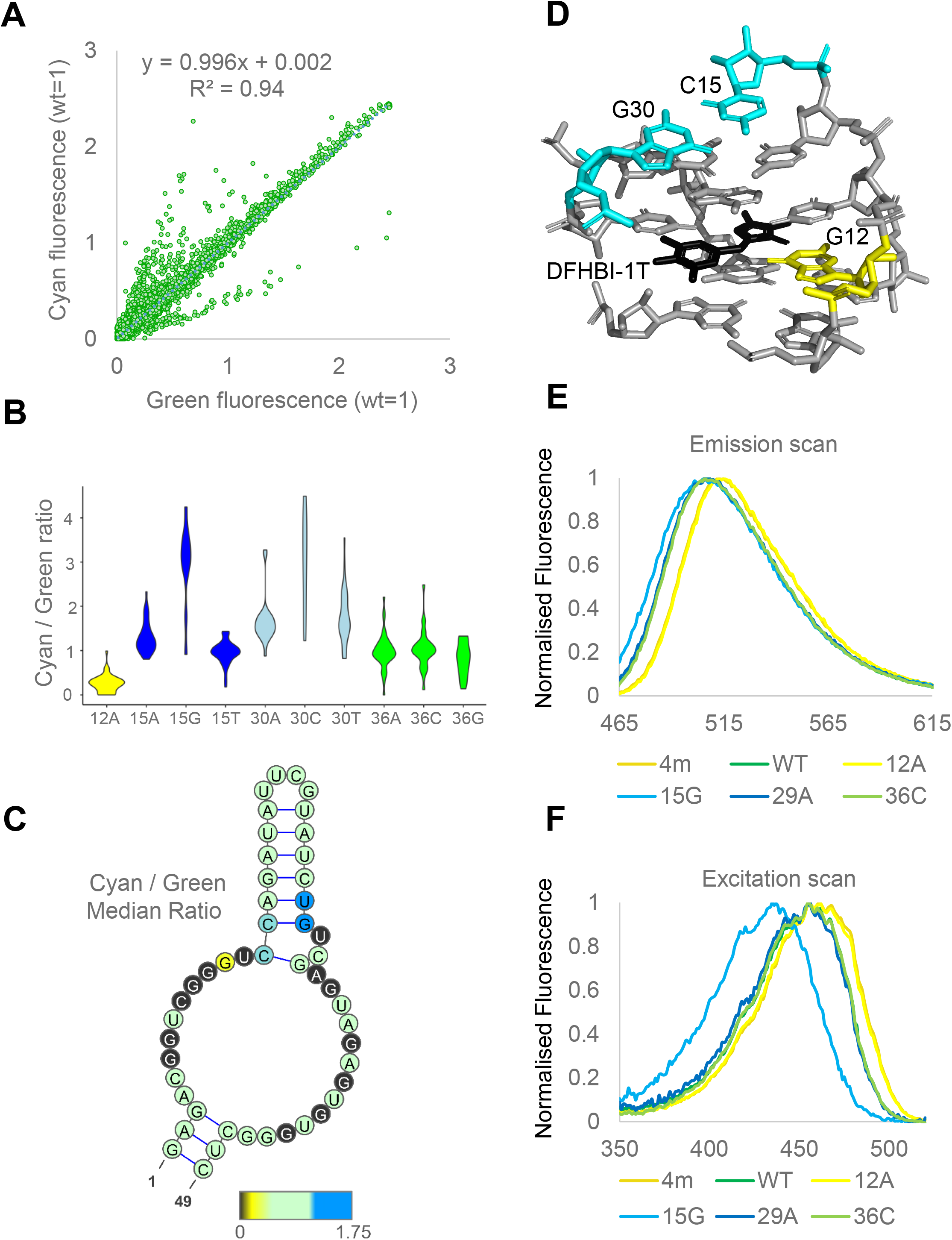
Comprehensive analysis of spectral tuning positions in Broccoli. (A) Fluorescence of Broccoli variants measured in the GFP channel (X axis) and CFP channel (Y axis). (B) Ratio of cyan to green fluorescence in single mutants and a subset of double mutants that comprise a mutation at the indicated position (see Methods). (C) Ratio of cyan to green fluorescence mapped onto the secondary structure. The colour of each position represents the median cyan/green ratio among single mutants and a subset of double mutants with a mutation at the focal position. (D) Positions with largest effects on fluorescence spectra mapped to 3D structure. DFHBI-1T is shown in green. (E-F) Emission (E) and excitation (F) spectra of selected variants measured in a plate reader.

To analyse the spectral shifts in more detail, we measured the fluorescence spectra of selected variants in a spectrofluorometer. The C15G mutation, which destabilised the stem above the fluorophore binding pocket, shifted the excitation maximum to 435 nm with almost no change to the emission spectrum (Fig. 4D). The G12A mutation, in the presence or absence of other mutations, caused a yellow-shift in both the emission and excitation spectra (Fig. 4D). In a previous study, trifluoromethyl groups in DFHBI derivatives changed fluorescence spectra by altering dipole moments across the fluorophore, and the G12A mutation presumably mimics this effect by disrupting the hydrogen bonding with the carbonyl oxygen of DFHBI-1T (Fig. 4E). Measurement of fluorescence spectra in solvents with different dielectric constants confirms that changes of polarity may induce blue and yellow shifts in DFHBI-1T (Fig. S12).

## Conclusion

A number of fluorescent RNA aptamers have been developed in the past decade (Chen et al., 2019; Dolgosheina et al., 2014; Filonov et al., 2014; Paige et al., 2011; Song et al., 2017; Steinmetzger et al., 2019) with emerging applications such as monitoring of transcription (Song et al., 2017), visualisation of genomic loci (Chen et al., 2019), and sensing of metabolites, signaling molecules, ions, and drugs ((Kellenberger et al., 2013; Paige et al., 2012), reviewed in (Su and Hammond, 2020)). *In vivo* applications of such aptamers require the optimization of molecular phenotypes, such as the fluorescence spectrum, brightness, affinity to fluorophore and other ligands, sensitivity to pH and metal ions, photobleaching, thermal stability and resistance to helicases and nucleases. Many of these properties can be adjusted by mutations in the RNA (Ageely et al., 2016; Filonov et al., 2014; Warner et al., 2014). However, currently there are no established methods to predict how individual mutations will affect most phenotypes of interest, and, as a result, aptamer construction is typically done by random mutagenesis followed by functional screening (Filonov et al., 2014; Han et al., 2013; Song et al., 2017; Strack et al., 2013). Our dataset provides a systematic overview of how mutations in the Broccoli RNA aptamer influence parameters relevant to in vivo function, paving the way to the development of predictive machine learning models that will allow the design of RNA molecules with required properties. The recent success of AlphaFold in the prediction of protein structures highlights the benefits of large datasets in understanding molecular function (AlQuraishi, 2019). We anticipate that our microarray platform, which uses microscopy equipment readily available in molecular biology labs, will facilitating both the development of RNA tools and a mechanistic understanding of their function.

## Acknowledgments

We thank Toby Hurd for the inspiration to start this project; Duncan Sproul for suggesting to use a microarray platform; members of the Advanced Imaging Facility in IGMM Central for help with microscopy; Hubert Czaja for help with computational analyses, Filip Stefaniak, Pritha Ghosh, Guido Sanguinetti and members of the Kudla laboratory for discussions. V.S.B. is a cross-disciplinary post-doctoral fellow supported by funding from the University of Edinburgh and Medical Research Council (MC_UU_00009/2). This work was supported by the Wellcome Trust (Fellowship 207507 to GK) and the Medical Research Council (grant MC_UU_00007/12 to GK).

## Author contributions

GK and OP conceived the work and designed the experiments and analyses. OP, GS, HI and MV performed experiments. OP, VSB, DAO, JMB, GW and GK performed computational analyses. GK wrote the paper.

## Materials and Methods

### DNA library design and amplification

We designed a pool of oligonucleotides composed as follows: truncated T7 polymerase promoter “ACGACTCACTATAGGGAGA” (19nt) – unique probe sequence (60nt) - “AAAAAAAAAAAAAAAA” (16nt) – upstream part of F30 “TTGCCATGTGTATGTGG” (17nt) – Broccoli variant (49nt) – downstream part of F30 “CCACATACTCTGATGATCCTTCGGGATCATTCATGGCAA” (39nt). As the oligo length was limited to 200 nucleotides, part of the T7 polymerase promoter was added during amplification of the library. The 60-mer probe sequences were derived from the GPL10787-9758.txt file containing information for Agilent SurePrint G3 Mouse GE 8×60K Microarray (Glass slide formatted with eight high-definition 60K arrays), that includes probes for mRNAs and lincRNAs. Sequences of Broccoli variants were generated by a custom script that changed the original nucleotide in each position of an input sequence to each of the 3 remaining nucleotides, producing 3 x “length of the sequence” of mutated variants. The first round was run on wild type Broccoli (“GAGACGGTCGGGTCCAGATATTCGTATCTGTCGAGTAGAGTGTGGGCTC”), to generate single mutants. This output was use as input for a second round – to generate double mutants.

A chemically synthesised single-stranded DNA library containing 1 fmol (~0.15 ng) of each oligonucleotide was purchased from Twist Bioscience. After optimising PCR conditions using a single variant of wild-type Broccoli oligo DNA, the whole library of Broccoli mutants was amplified. 50 ng of the oligo DNA library was used for PCR amplification with Taq DNA polymerase (Invitrogen, Cat No. 10342-020), forward primer (AAAATTGCCATGAATGATCCCGA), reverse primer (TAATACGACTCACTATAGGGAGA) using the optimised protocol. PCR mix was made in a final volume of 50 μl (Template library DNA – 50 ng in 10μl; forward primer – 0.7 μM final conc.; reverse primer – 0.7 μM final conc.; 10× PCR Buffer w/o Mg – 1× final conc.; MgCl_2_ – 2 mM final conc.; dNTP’s – 0.25 mM each final conc.; Taq DNA polymerase – 10U) and the PCR thermocycler program was set as follows. Initial denaturation at 95°C for 3 min. was followed by 9 – 12 three-step cycles of: denaturation at 95°C for 15 s.; annealling at 55°C for 30 s.; extension at 68°C for 90 s. The PCR product was purified with MinElute PCR Purification Kit (Qiagen, Cat No. 28004) following manufacturer’s protocol using a double elution with 20 μl of Elution Buffer each. The final DNA concentration was determined using NanoDrop 8000 spectrophotometer (ThermoFisher, Cat No. ND-8000-GL).

### In vitro transcription

All of the purified DNA product from the library amplification PCR was used as a template for *in vitro* transcription reactions using MEGAshortscript T7 Transcription Kit (ThermoFisher, Ambion, Cat No. AM1354). The transcription reactions were performed following the manufacturer’s protocol and using 200 nM final concentration of template DNA (~550 ng of template DNA per 20 μl reaction). Reactions were incubated at 37°C overnight (18-24 h) in ThermoMixer F1.5 (Eppendorf, Cat No. 5384000039) with ThermoTop (Eppendorf, Cat No. 5308000003) followed by TURBO DNase treatment at 37°C for 20 min. The RNA product from each reaction was purified with RNeasy MinElute Cleanup Kit (Qiagen, Cat No. 74204) following a manufacturer’s protocol using a double elution with 40 μl of RNase-free water each. The final purified samples were mixed together and the RNA concentration was determined using NanoDrop 8000 spectrophotometer. A sample of the final RNA product was run on an agarose gel to confirm the correct size and check for RNA quality and possible degradation.

### RNA labelling with Cy3

Ten percent of the final RNA amount was used for chemical labelling with Cy3 fluorescent dye using Arcturus Turbo Labeling Cy3 Kit (ThermoFisher, Applied Biosystems, Cat No. KIT0609). The chemical labelling procedure and subsequent removal of free Cy3 dye was performed following manufacturer’s protocol with 15 μg of RNA per reaction. The labelled and purified RNA from all labelling reactions was mixed together and the final RNA and Cy3 concentrations were determined using NanoDrop 8000 spectrophotometer.

### Hybridisation to microarrays

The complete RNA library was hybridised to SurePrint G3 Mouse GE 8×60K Microarrays (Agilent, Cat No. G4852A) using a partially modified version of manufacturer’s protocol as described in Version 6.9.1 of ‘One-Color Microarray-Based Gene Expression Analysis Protocol’. The total amount of RNA used for hybridisation on each microarray was between 20 μg and 45 μg and contained between 1% and 7% of Cy3 labelled RNA. The most robust imaging data were obtained after hybridisation with 40 μg of total RNA containing 2.5% of Cy3 labelled RNA (39 μg of unlabelled RNA mixed with 1 μg of Cy3 labelled RNA). The hybridisation mix was prepared by mixing the specified amounts of RNA with 10× Gene Expression Blocking Agent (1× final conc.) and 2× Hi-RPM Hybridization Buffer (1× final conc.) (Agilent, Cat No. 5188-5242) in a final volume of 42 μl. The RNA Fragmentation Buffer was not used and the incubation at 60°C for RNA fragmentation was not performed. 40 μl of the hybridisation mix was transferred into each of eight wells on a hybridisation gasket slide (Agilent, G2534-60015) during hybridisation assembly using the Agilent Microarray Hybridisation Chamber Kit (Agilent, Cat No. G2534A) and following the standard protocol. Assembled chambers were incubated in a rotating hybridisation oven at 65°C for 18 hours. After hybridisation microarray slides were washed with Gene Expression Wash Buffer 1 and Gene Expression Wash Buffer 2 (Agilent, Cat No. 5188-5327) with added Triton X-102 (0.005% final conc.) following the standard manufacturer’s protocol.

### Imaging buffer composition

Microarray imaging was performed in an imaging buffer solution containing fluorophore DFHBI-1T ((Z)-4-(3,5-difluoro-4-hydroxybenzylidene)-2-methyl-1-(2,2,2-trifluoroethyl)-1Himidazol-5(4 H)-one) (excitation = 472 nm, emission = 507 nm)) from Lucerna Technologies Cat. no. 410, MgCl_2_, KCl and HEPES. A series of imaging buffers were used for imaging in concentration gradients of fluorophore and salts as well as pH and temperature gradients. The different components were tested in the following concentration gradients. DFHBI-1T concentration gradient (μM): 0.1; 0.25; 0.5; 1; 2; 5; 10; 20; 40; 80; the standard concentration for other gradients was 10 μM. MgCl_2_ concentration gradient (mM): 0; 0.05; 0.1; 0.2; 0.5; 1; 3; 5; 10; 30; concentrations used for other gradients were 1 mM, 5 mM and 10 mM. KCl concentration gradient (mM): 2; 5; 10; 20; 40; 80; 140; 200; the standard concentration used for other gradients was 140 mM. A pH gradient was made using 20 mM HEPES with pH – 5; 5.5; 6; 7; 8; 9; pH values used for other gradients were 5.5 and 7. A temperature gradient was generated by adjusting the temperature in the microscopy room using air conditioning and by using heated chamber with a thermostat fitted on the microscope. The temperatures tested were: 17°C; 23°C (using A/C setting); 30°C; 37°C and 42°C (using heated chamber). The standard temperature for other gradients was 23°C.

### Imaging chamber assembly

A major technical challenge was to keep microarrays in a constant concentration of imaging buffer components, avoiding any evaporation throughout the whole duration of an imaging session that lasted about one hour. For that purpose specially designed single-use imaging chambers were made using 18 mm × 18 mm cover glasses #1 (VWR, Cat No. 631-1331) and silicon seals cut out of ultra thin silicone film with 0.3 mm thickness (Silex, Ultra Thin Silicone Film – 0.3 mm – 30 Shore). Silicone seals were cut out using precision CNC cutting machine with inner dimensions of 10 mm × 13.5 mm and 1.5 mm width at each side. First the cover glasses were gently but thoroughly washed with 80% ethanol and lens cleaning tissues, then gently dried with lens cleaning tissues and dusted with manual air duster. Using forceps, the silicone seals were very carefully placed with the more sticky side facing the cleaned cover glasses leaving about 5 mm space on one side of the cover glasses for handling. Such prepared imaging chambers were stored in an empty cover glass box held by sponge at the sides to prevent touching each other. Imaging chambers were handled with care to avoid ever touching the imaging area and were air dusted again before use. Just before starting an imaging session 25 μl of imaging buffer was carefully placed in the centre of the microarray and the imaging chamber was precisely positioned on this microarray using forceps and was gently pushed along the edges using a tip of the forceps to seal the chamber. Any droplets of the imaging buffer outside of the chamber were immediately dried using a corner of a tissue paper. A clean hybridisation gasket slide and an old microarray slide stained with ink were placed precisely underneath the fresh microarray slide to indicate the position of the transparent microarrays and guide placing imaging buffer and chamber on the microarray. A well positioned flat silicone seal would prevent any evaporation of the buffer from the imaging chamber. After the imaging was complete a buffer change was performed by swiftly and carefully detaching the imaging chamber from the microarray slide using forceps and then briefly drying the microarray slide by gently tapping one side on the bench with clean tissue paper underneath to absorb the buffer. Another sample of imaging buffer and fresh imaging chamber were quickly placed on the same microarray. The buffer change had to be performed relatively quickly to avoid a complete drying out of the microarray as this would often lead to deterioration of the spots on the microarray. Usually three to four buffer changes were made but sometimes even up to ten buffer changes were performed on one microarray still giving good quality data, however this varied depending on the hybridisation and imaging conditions and had to be assessed individually for each microarray.

### Fluorescence microscopy

Microarrays with assembled imaging chambers were imaged through the cover glass and the imaging buffer using Zeiss ‘Observer.Z1’ inverted epifluorescence microscope with dry 10x/0.45 Plan-Apochromat objective (Zeiss, Cat No. 420640-9900-000) and ‘Retiga 6000’ monochromatic 14 bit cooled CCD camera with 2750×2200 pixels and 4.54 μm pixel size (QImaging, 01-RET-6000-R-M-14-C). The microscope was equipped with an LED light source and the following filters were used for each channel: green (GFP) channel excitation – 474/27, emission – 520/35; red (RFP) channel excitation – 554/23, emission – 609/54; cyan (CFP) channel excitation – 434/17, emission – 479/40. The microscope was controlled using the Micro-Manager software package (Edelstein et al., 2010).

The imaging was performed with the following microscope and camera settings: camera gain = 2; binning = 4; camera ROI after binning = 585×552 pixels; GFP exposure = 2000 ms; RFP exposure = 700 – 900 ms; CFP exposure = 1100 ms; light source intensity = 100% for all channels; Z stack = 3 – 4 with 0.015 μm steps. An autofocus ‘OughtaFocus’ algorithm was used with the following settings: SearchRange_um = 50; Tolerance_um = 1; CropFactor = 1; Exposure = 400 ms; ShowImages = Yes; Maximize = Sharp edges; Channel = RFP. Imaging of one microarray required 204 tiles using a grid of 17×12 tiles with 40% overlap between the tiles. Each tile was imaged 3 or 4 times in each channel as separate Z stacks with virtually identical focal plane as the 0.015 μm step between the stacks was below the stage motor resolution. These separate images for each tile were later averaged during processing to reduce the background noise.

### Feature extraction

The individual image files were combined into a single image of the entire chip using a bespoke LabView (National Instruments) program which corrected for illumination heterogeneity before merging the images together. Areas of overlap were added together using linear opacity ramps to avoid hard edges between images. Multiple acquisitions of the same colour and settings were averaged to improve signal to noise ratio.

The arrangement of the array of features was identified by matching the fixed patterns of spots in the four corners of the array using the green channel, and this information was used to identify the approximate position and index number of each spot. These positions were refined by searching for a circular pattern of bright pixels starting from the initial approximate location. The average pixel value was extracted for each feature in all the colours recorded and saved. Each feature was then classified by its index number into groups for easier processing.

### Calculating the fluorescence of Broccoli variants

The microarray contained 40371 spots expected to bind Broccoli variants (78 spots for wild-type Broccoli, 8100 single or double mutants with 4 spots per variant, and 2631 variants with 3 spots per variant), as well as 9925 “empty spots” where nothing was designed to hybridise, but where we could still read signal. We concluded that the signal might be coming from non-specific cross-hybridisation so the average signal of these empty spots was subtracted from all values in each channel. To account for the differences in the amount of Cy3-labeled RNA used in each experiment, we divided the red signal of each spot by the mean red signal of all spots on a given array. Spots with red signal less than 0.2 (i.e. less than 20% of average red signal on the array) were excluded from further analysis. All spots with negative green signal had the green signal set to 0 and the normalized Broccoli fluorescence in each spot (reported in the master data table), was calculated as (green signal) divided by (red signal +1). For downstream analyses, we then calculated the median fluorescence across spots and across experiments performed in the same condition.

### Modelling of epistasis

The global epistasis model was based on data from eight replicate experiments performed in a buffer that contained 10 mM Mg^2+^, 10 uM DFHBI-1T, 140 mM K^+^, pH=5.5, T=23°C. Prior to modelling, we normalized green fluorescence of each spot by (1) dividing the green fluorescence by (red fluorescence+1), and (2) dividing the result by the median green fluorescence in each experiment.

The global epistasis model estimates the fluorescence of double mutants as a function of the fluorescence of the two single mutations found in the double mutant, without taking into account the exact identity of these single mutations. Thus, for each double mutant mut_ij_, the fluorescence is calculated as:

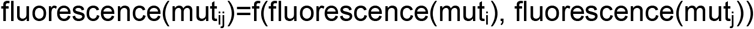

In the equation above, mut_i_ and mut_j_ are single mutants; fluorescence(mut_i_) and fluorescence(mut_j_) are estimated fluorescence values of these single mutants (see below), and f is a locally weighted polynomial regression (LOESS) function, which ensures a smooth dependence between the inputs (fluorescence of single mutants) and the output (fluorescence of double mutants). To estimate the fluorescence of single mutants for use in the equation above, we used the mean fluorescence of spots representing the single mutants themselves, and spots representing double mutants that included the focal mutation and a mutation in one of the loop positions (20, 21, 22, 23, 24, 25), which had little effect on fluorescence. We found that using a subset of double mutants in the estimation of single mutation effects increased the amount of available data, reduced noise, and did not bias the results. We calculated global epistasis using the “loess” function in R. To calculate local epistasis values for all double mutants, we subtracted the measured fluorescence of each double mutant from the fluorescence predicted using the global epistasis model described above.

### Training of machine learning models

We considered all pairs of positions (i,j) and labelled each pair as either paired (10) or non-paired (980) based on the secondary structure model of wild-type Broccoli. We excluded pairs in which |i-j| < 4 because such pairs are not commonly found in RNA structures. We trained an ensemble of Support Vector Machine (SVM) classifiers on the labelled data, using the scikit-learn library in Python with a 70:30 split between training and test datasets. For model training, we employed as features the mean local epistasis signals for the A:U, G:C, and G:U pairs:

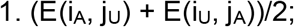

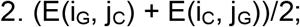

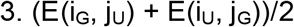

where E(i_X_, j_Y_) is the value of local epistasis of a double mutant with mutation X in position i, and mutation Y in position j. We randomly shuffled the data a total of 1000 times, from where we built an ensemble of 1000 classifiers and computed the frequency of each possible pair being called as paired in the test data.

### Curve fitting

We used the open source tool gnuplot to calculate the dissociation constant (Kd) and maximum fluorescence (F_max) of the Broccoli-DFHBI-1T complex by fitting curves described by a first order Hill equation (Hill coefficient=1) to data from DFHBI-1T gradient experiments:

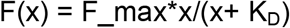

In this equation, x is the concentration of DFHBI-1T (in μM units) and F(x) is the Broccoli fluorescence measured in the given concentration of DFHBI-1T. We assumed that the fluorescence without fluorophore was equal to 0: F(0)=0. We excluded variants with low fluorescence (less than 8% of wild-type fluorescence with the highest DFHBI-1T concentration), and variants for which Kd or Fmax could not be reliably estimated (Kd(DFHBI-1T) > Fmax(DFHBI-1T) or Kd(DFHBI-1T) < 0.5 μM or Kd(DFHBI-1T) > 40 μM), leaving ~3,500 variants with Kd and F_max estimates. Calculations of EC50(Mg^2+^) and EC50(K^+^), defined as the concentrations of magnesium and potassium ions that elicited half of the maximum fluorescence in magnesium and potassium gradient experiments, were performed in a similar way. We excluded variants for which the estimated EC50 value was greater than the respective F_max, or where the estimated EC50 was negative, or F_max was greater than 1000 fluorescence units.

### Fluorescence measurements of individual variants

To confirm and further explore the fluorescence properties of chosen Broccoli mutants we ordered 4 nmole Ultramer® DNA Oligo from IDT (Integrated DNA Technologies), encoding each variant fused with the F30 scaffold under a T7 promoter. After annealing with an antisense oligo (same as for transcribing the library) in the promoter region, the ssDNA served as a template for production of RNA with MEGAshortscript™ High Yield Transcription Kit (Thermo Fisher Scientific) following the manual, with the incubation in 37°C time extended to 48h. The RNA product from each reaction was purified with RNeasy MinElute Cleanup Kit (Qiagen, Cat No. 74204) following the manufacturer’s protocol. To perform low-throughput measurements of fluorescence in an automated plate reader (Tecan Infinite M200 Pro), we used 0.5 μg of RNA in 50 μl of buffer. Each sample was measured in 4 repeats. The buffer composition was as follows:

- KCl 100mM; MgCl_2_ 10mM; DFHBI-1T varied; RNA 0.5μM; pH 5.5 for DFHBI dependence,
- KCl 100mM; MgCl_2_ varied; DFHBI-1T 40μM; RNA 0.5μM; pH 7.4 for magnesium dependence,
- KCl varied; MgCl_2_ 1mM; DFHBI-1T 40μM; RNA 0.5μM; pH 7.4 for potassium dependence
- and KCl 100mM; MgCl2 10mM; DFHBI-1T 40μM; RNA 0.5μM; pH 7.4 for excitation and emission spectra.

### Measuring of fluorescence spectra in solvents with different dielectric constants

The RNA was aliquoted and dissolved in DMSO (10 mM: 1 mg per 0.312 mL). Solutions were prepared in different solvents (final RNA concentration: 100 uM, maximum amount of DMSO: 1%). Emission spectra were measured in Biotek spectrophotometer (excitation at 430 nm, emission measured from 460 nm to 600 nm (1 nm slit) at 27.5C (gain 120). Experiments were performed in duplicate.

---

**Figure S1.**
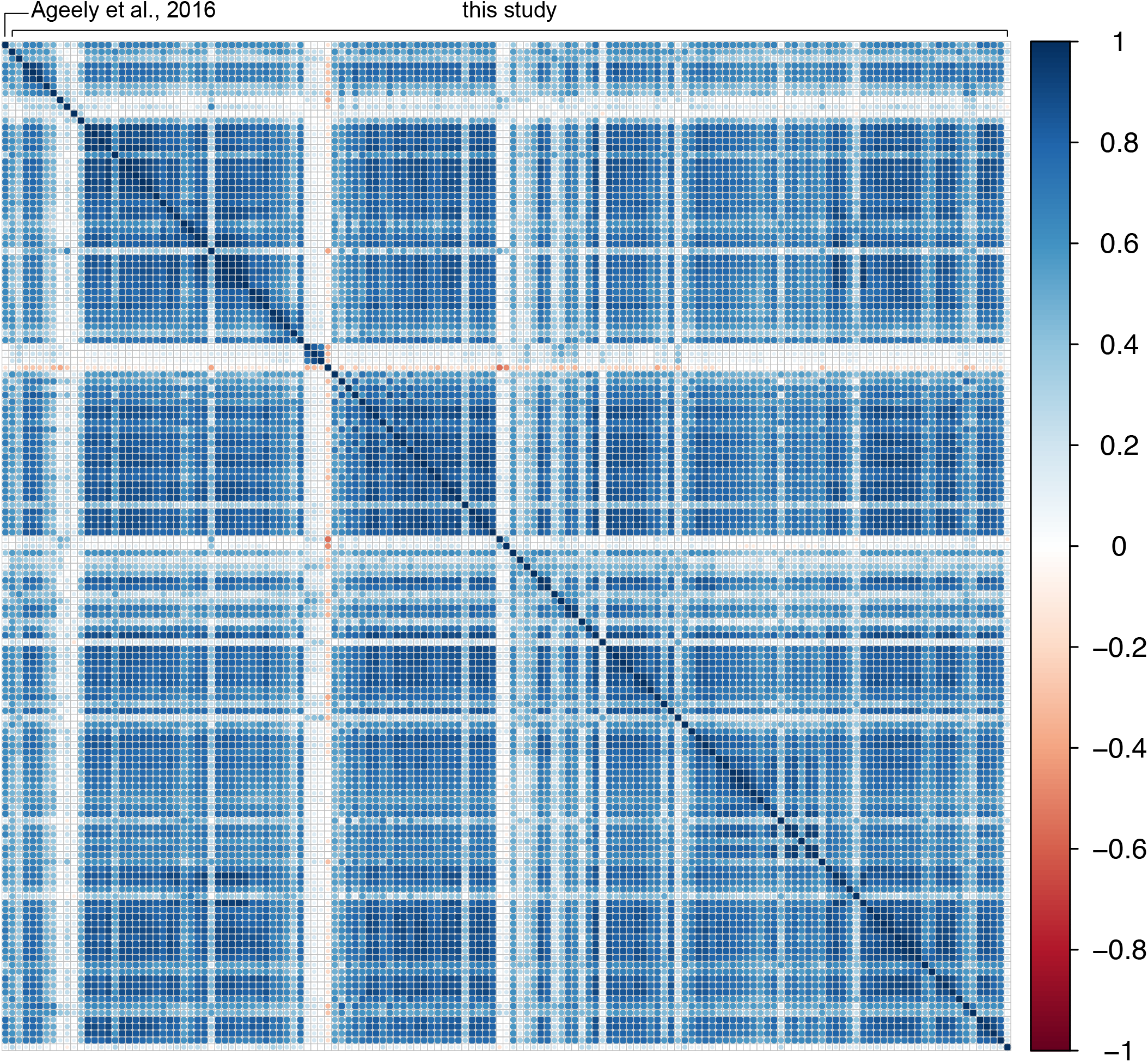
Reproducibility of fluorescence measurements. (A) Spearman correlation matrix of green fluorescence of Broccoli single mutants (N=147) in (Ageely et al., 2016) and in 146 experiments from the present study.

**Figure S2.**
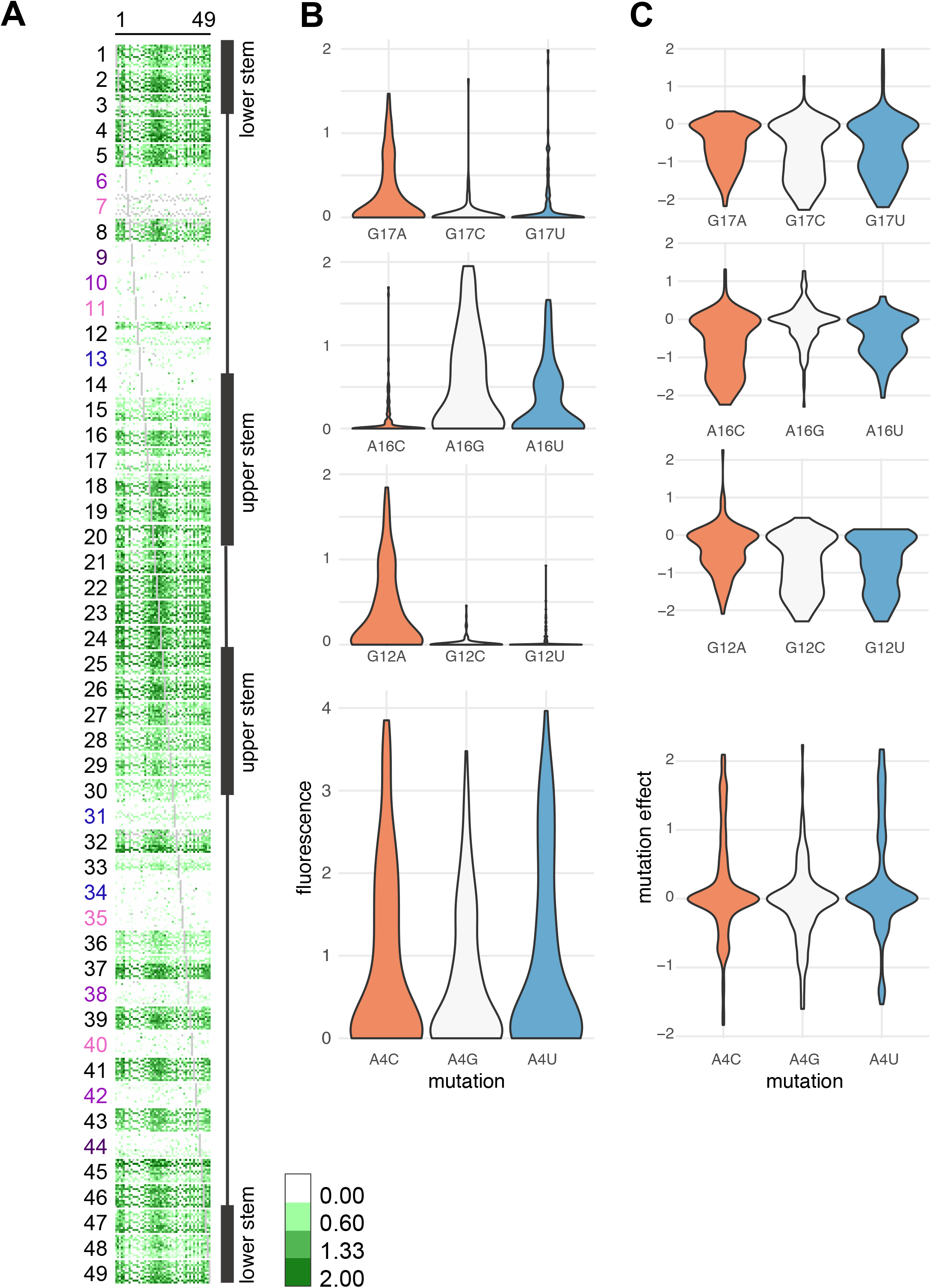
Single mutation effects depend on the genetic background. (A) Fluorescence of every possible mutation (X axis) in all 147 (3 × 49) single-mutation backgrounds (Y axis). Other details as in Figure 1C. Experimental conditions: 10 mM Mg^2+^, 10 uM DFHBI-1T, 140 mM K^+^, pH=5.5, T=23°C. (B) Distribution of fluorescence of all single and double mutants that contain the indicated mutation. (C) Distribution of effects of selected mutations across all genetic backgrounds. In this analysis, the effect of mutation i is calculated is calculated as effect(mut_i_) = (fluorescence(mut_ij_) - fluorescence(mut_j_)), where mut_ij_ is the double mutant with mutations i and j, and mut_i_ and mut_j_ are single mutants with mutations i or j.

**Figure S3.**
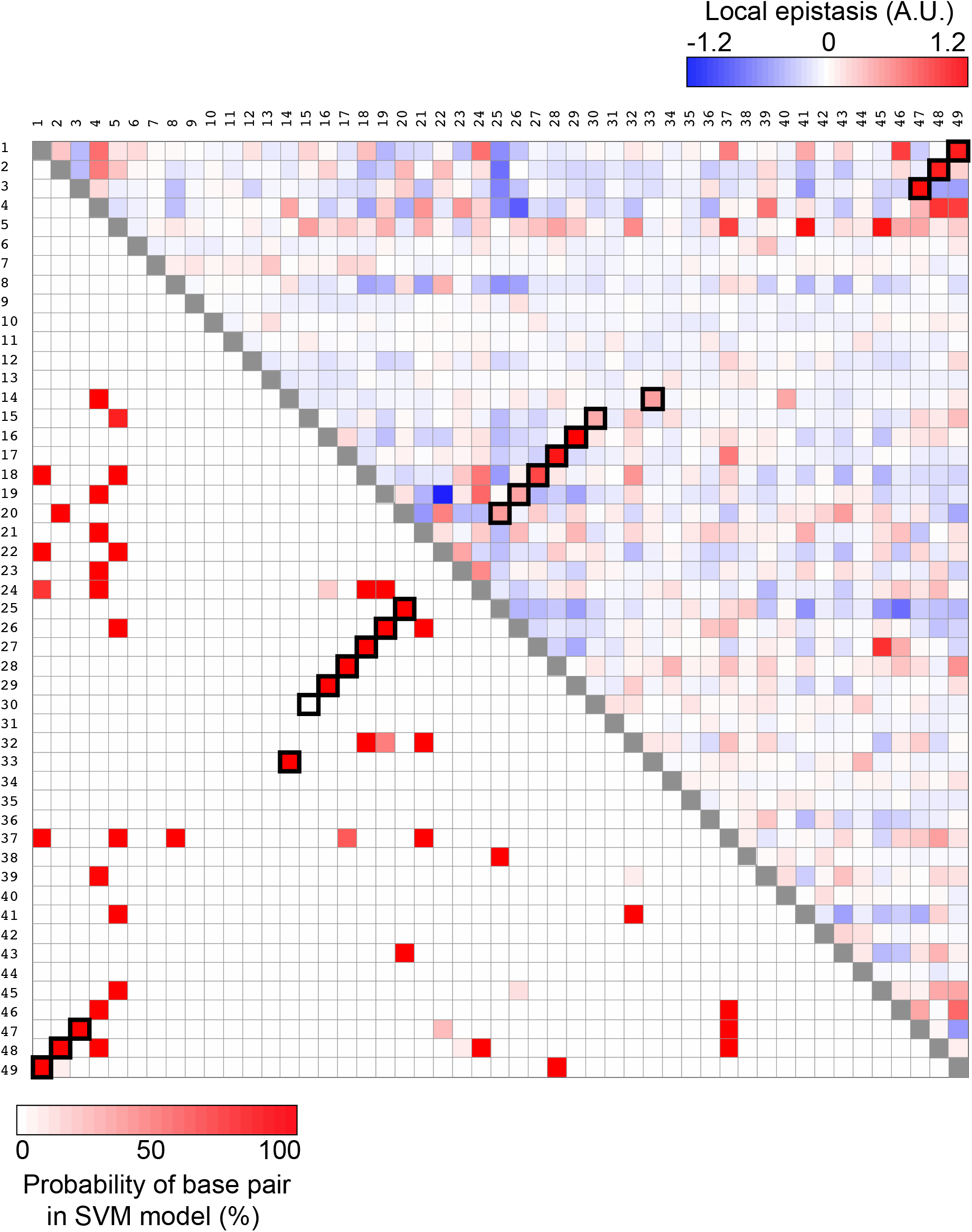
Matrix of local epistasis and predicted base pairing. (Top right) Mean local epistasis estimated for all pairs of positions as the difference between measured fluorescence, and fluorescence predicted with the LOESS model of global epistasis. Red colour indicates positive epistasis and blue, negative epistasis. (Bottom left) Probability for pairs of nucleotides to be identified as base paired by the SVM model. Saturated colour indicates higher probability. Base pairs found in the 3D structural model are shown by black square outlines.

**Figure S4.**
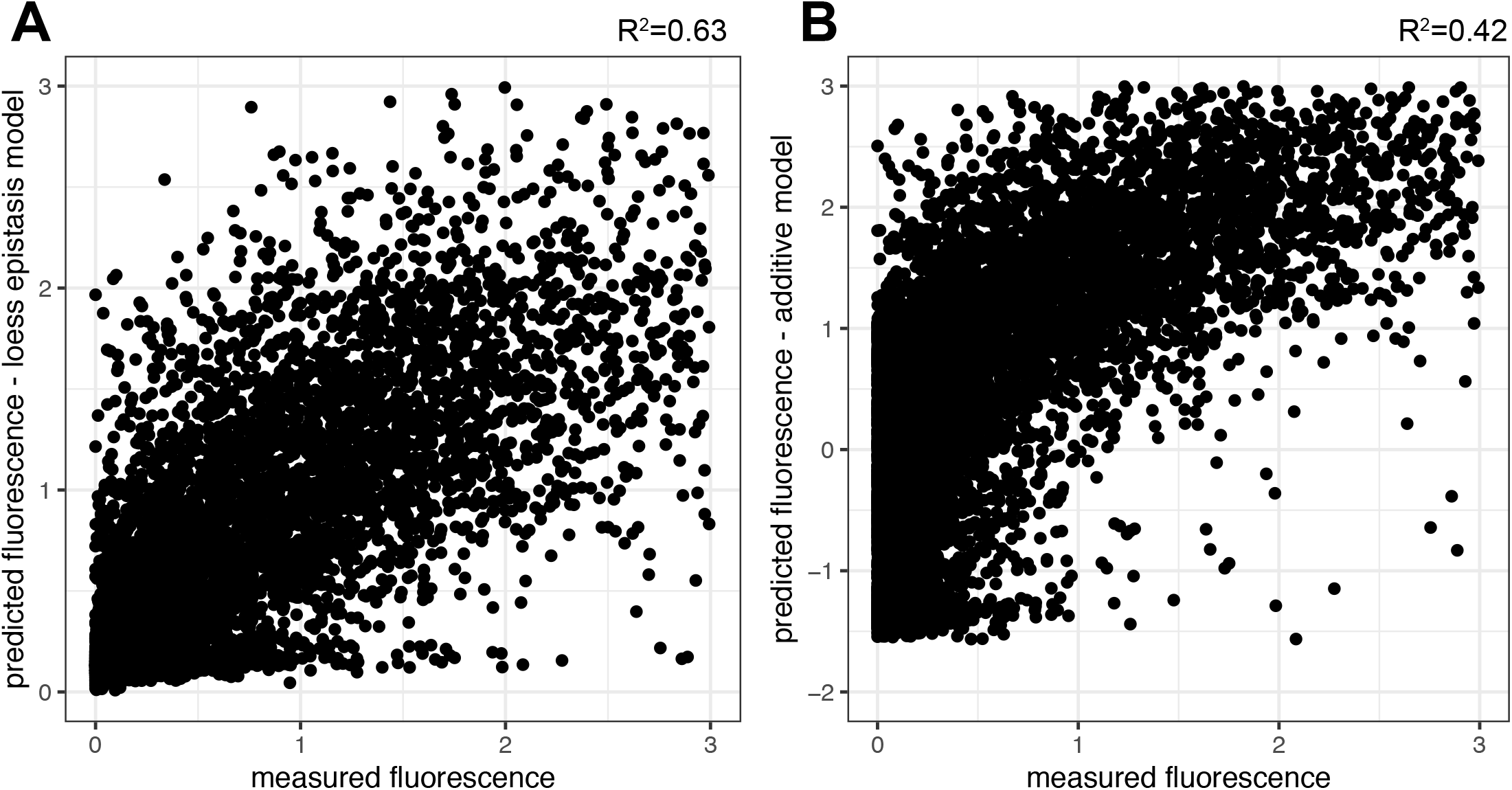
Comparison of measured and predicted fluorescence of double mutants. Fluorescence of double mutants was predicted from the fluorescence of single mutants using a loess model (A), or an additive model (B).

**Figure S5.**
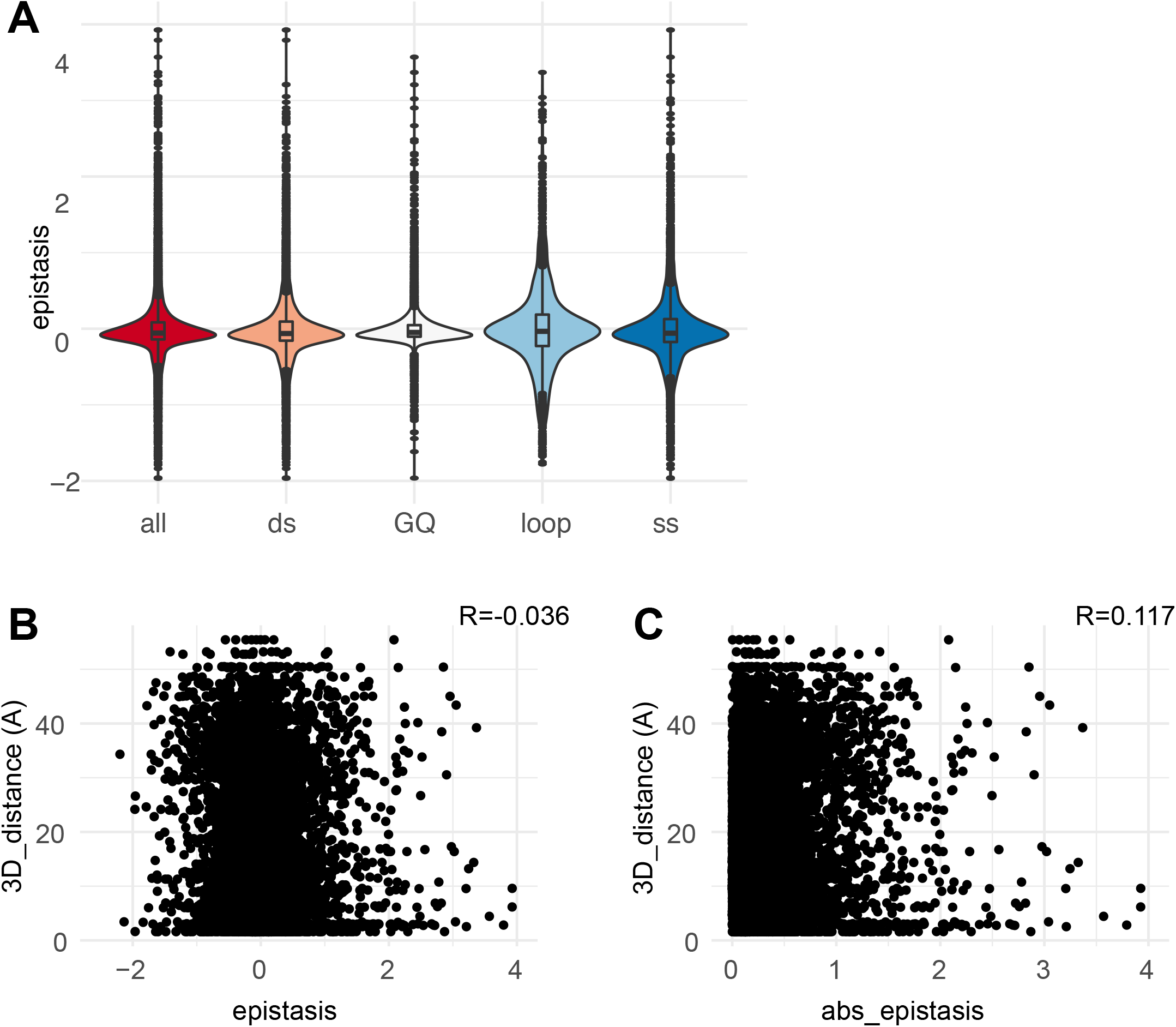
Structural correlates of local epistasis. (A) The distribution of local epistasis coefficients across structural categories. all, all sites; ds, double-stranded positions; GQ, G-quadruplexes and base triplets; loop, apical loop; ss, single-stranded positions. (B,C) Correlation of epistasis (B), or absolute value of epistasis (C) with distance between pairs of nucleotides in the 3D model.

**Figure S6.**
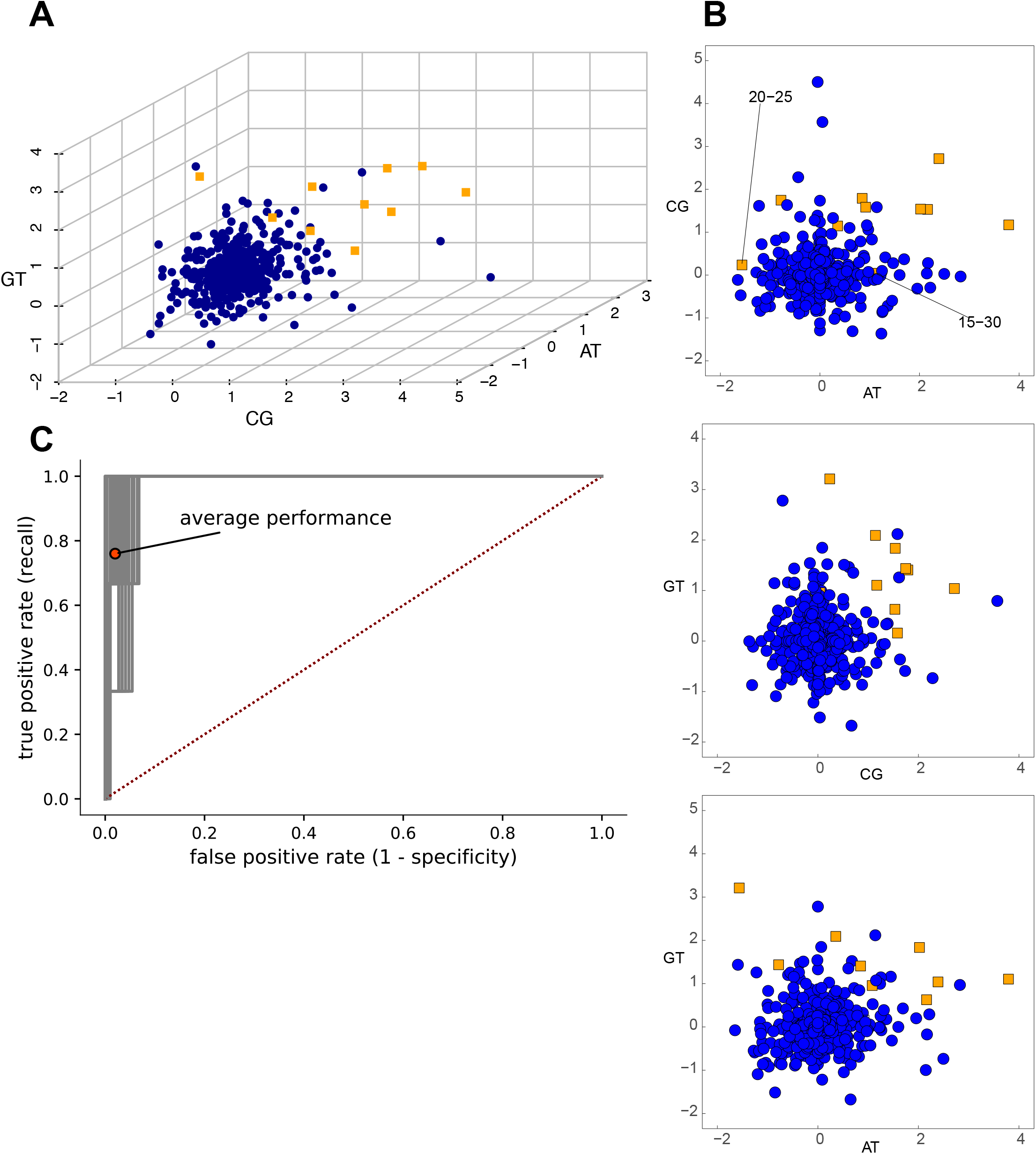
Prediction of RNA structure from epistasis signals. (A-B) Three-dimensional feature space for paired (orange) and non-paired (blue) bases of the wild-type Broccoli structure. The chosen features are the mean local epistasis signals for A:U, G:C, and G:U pairs (Methods), and were chosen on the basis of their ability to discriminate between paired and non-paired bases. The separation between both classes suggests the use of a support vector machine (SVM) as a classifier algorithm. The two pairs that were not efficiently predicted (pairs 15-30 and 20-25) are highligted. (C) Receiver operating characteristic (ROC) curve of the SVM trained on paired and non-paired bases of wild-type Broccoli. Shown are 50 ROC curves sampled from an ensemble of 1000 SVMs. The average recall and specificity are highlighted.

**Figure S7.**
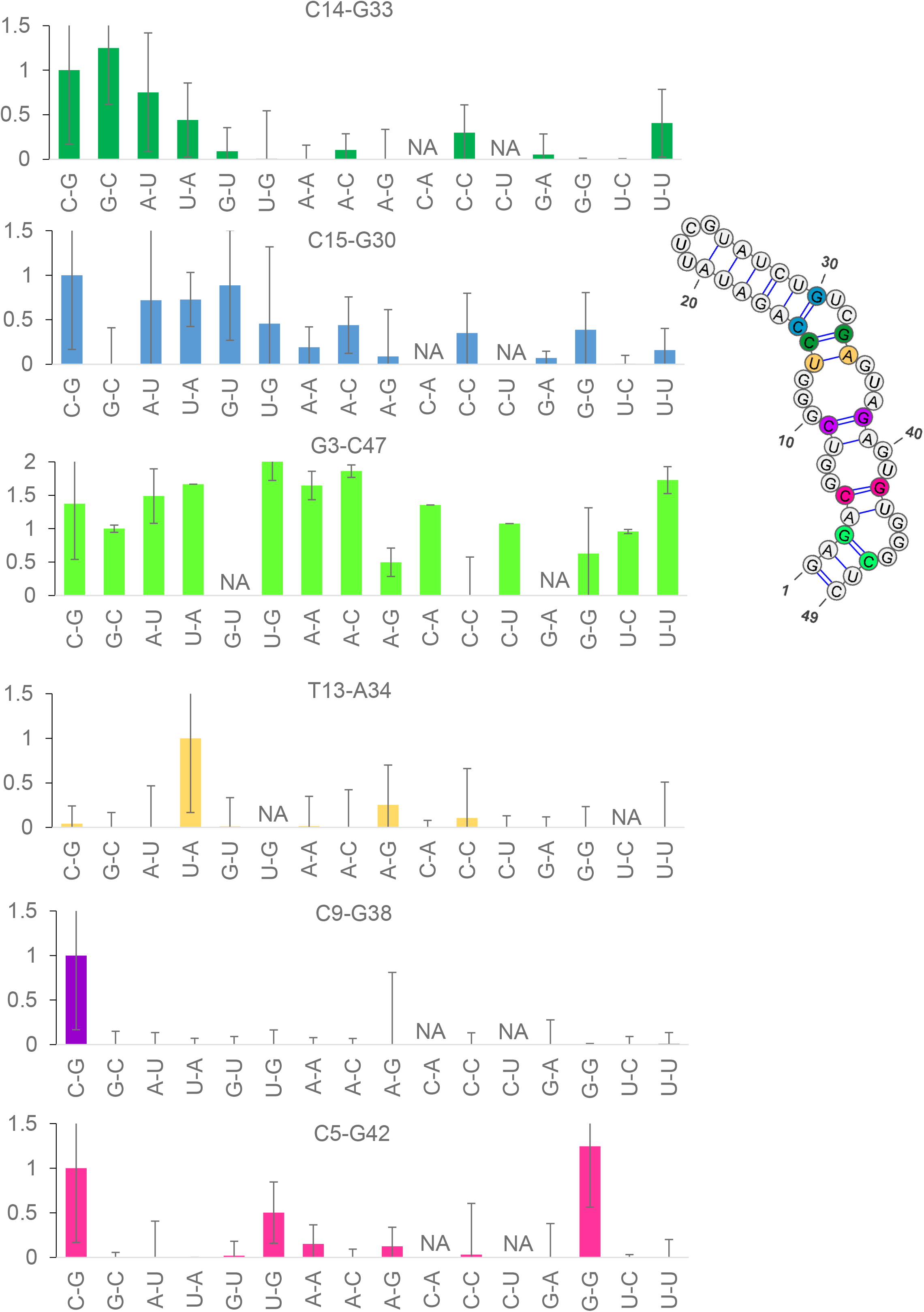
Compensatory mutations in base paired positions. Fluorescence of wild-type, single, and double-mutants in known base-paired positions (14:33, 15:30, 3:47), and in positions predicted as base-paired in the minimum free energy model, but not in the 3D structural model (13:34, 9:38, 5:42). NA, data not available.

**Figure S8.**
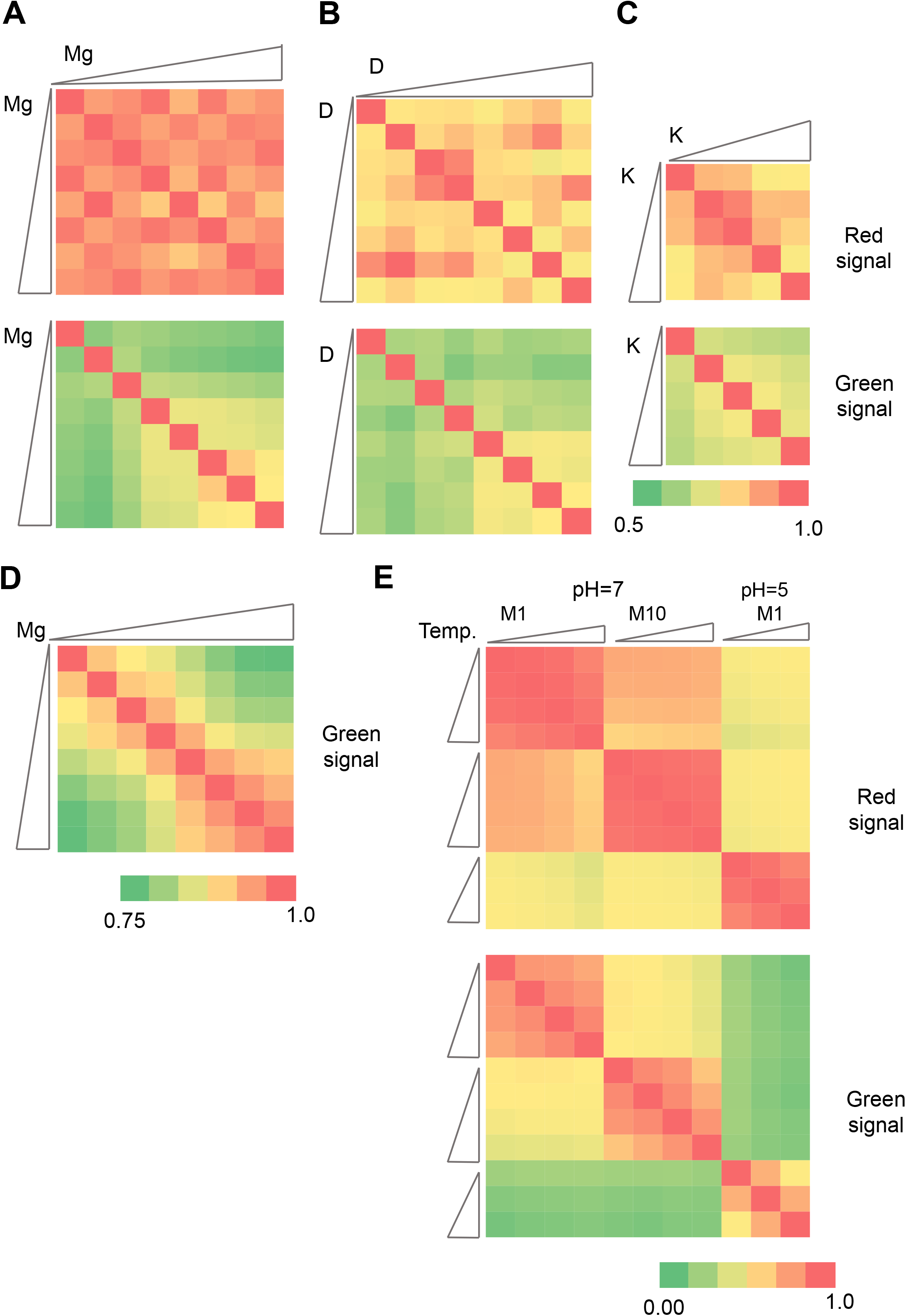
Comparison of fluorescence patterns across experimental conditions. (A-C) Correlations of red (top) and green (bottom) fluorescence of individual spots between experiments performed in increasing magnesium (A), DFHBI-1T (B) and potassium (C) concentrations. The exact composition of buffers is listed in the “Imaging buffer composition” section of the Methods chapter. (D) Correlations of green fluorescence of individual spots between experiments performed in increasing magnesium concentrations at pH=7. (E) Correlation of red (top) and green (bottom) fluorescence of individual spots between experiments performed in increasing temperature (indicated by triangles at the left and at the top) with varying pH and magnesium concentrations.

**Figure S9.**
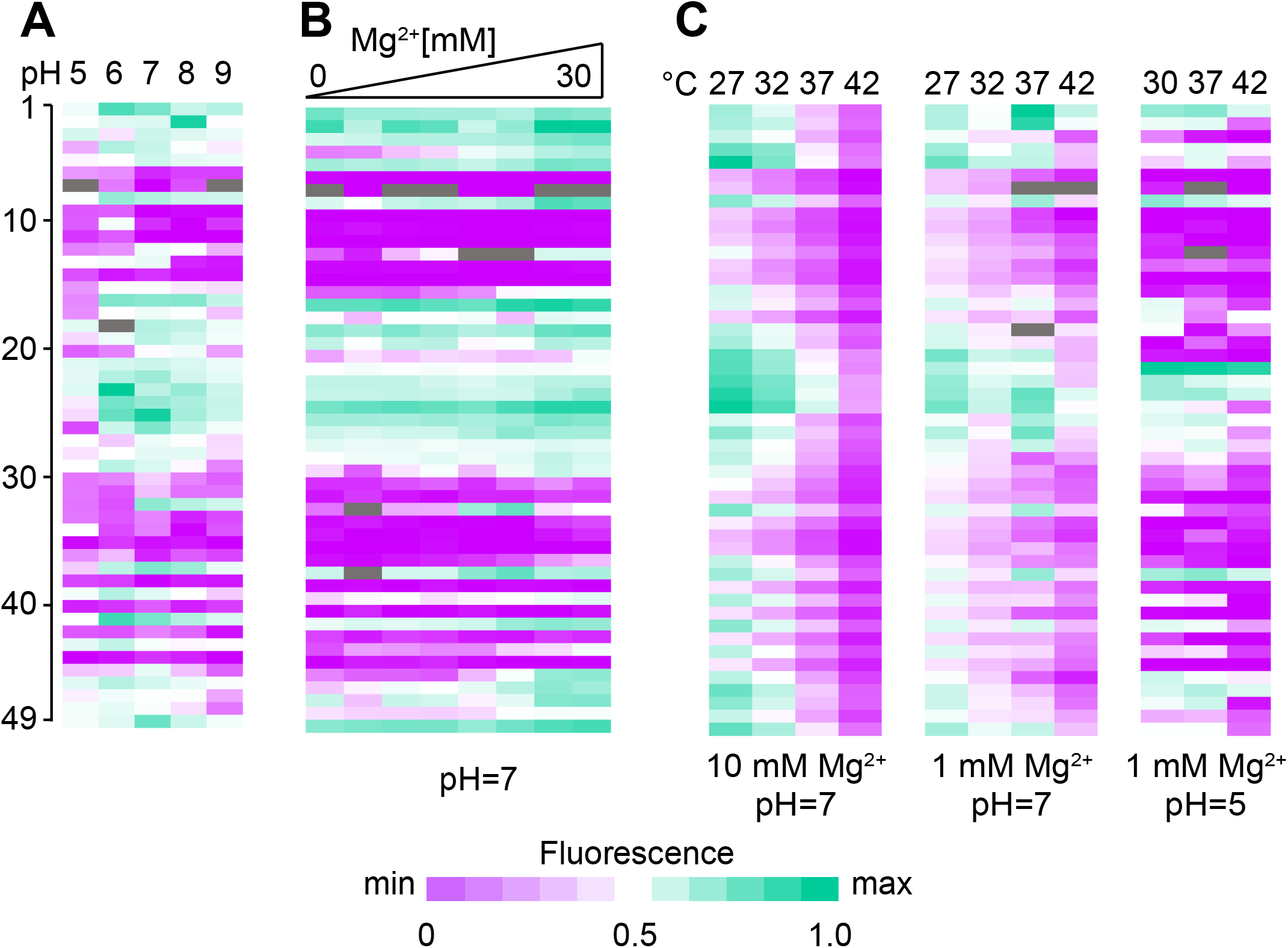
Effects of environmental conditions on fluorescence. (A) Fluorescence of Broccoli mutants as a function of the mutated position (Y axis) and pH (X axis). Each data point represents the median fluorescence of single mutants and a subset of double mutants that included a mutation at the indicated position (see Methods). (B) Fluorescence of Broccoli mutants as a function of the mutated position (Y axis) and Mg^2+^ concentration at pH=7 (X axis). (C) Fluorescence as a function of temperature measured at different pH and Mg^2+^ concentrations.

**Figure S10.**
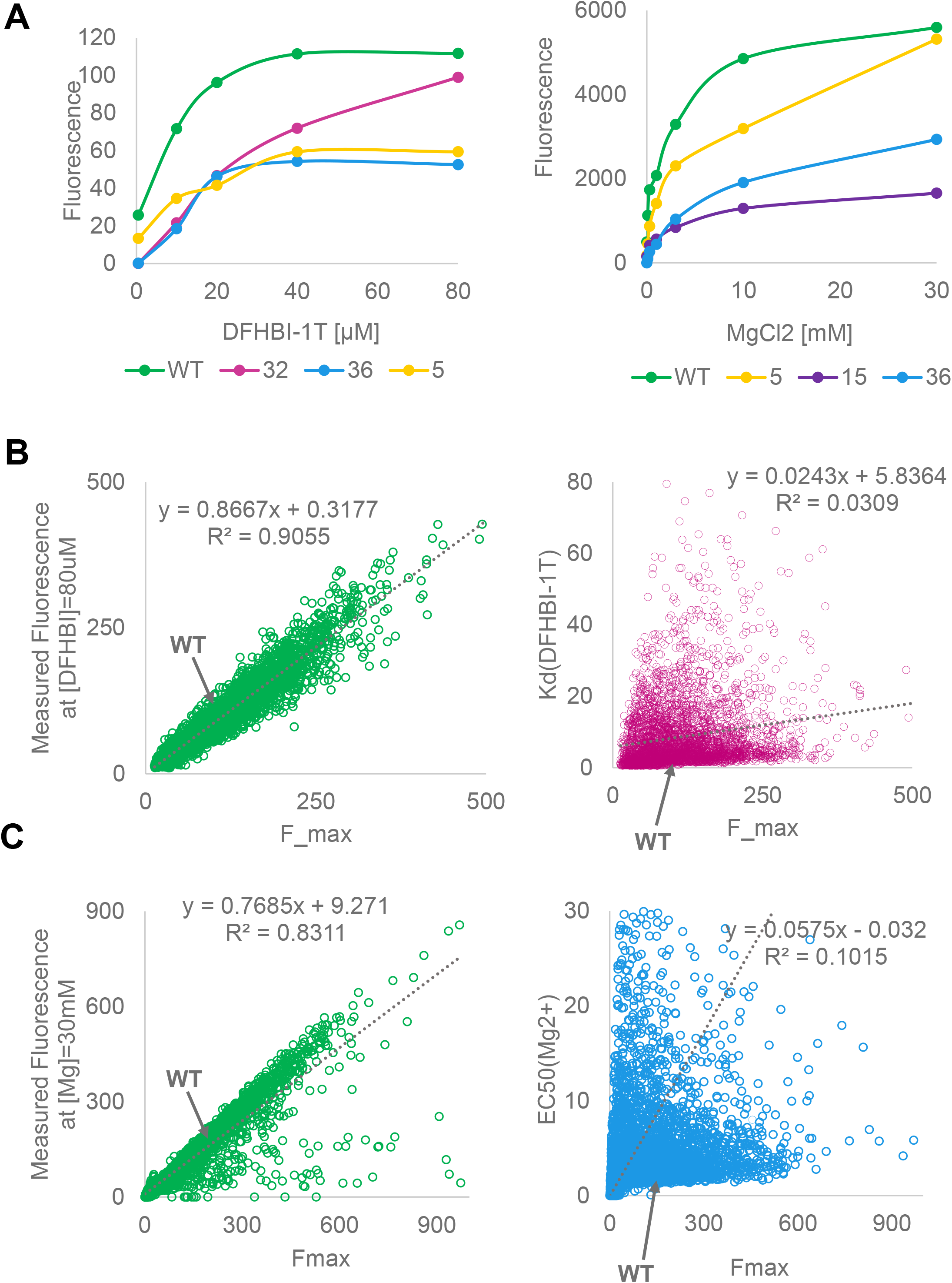
Microarray analysis of dissociation rate constants. (A) Median fluorescence of variants with single mutations at the indicated positions in increasing concentrations of DFHBI-1T and magnesium. (B) Correlations between measured fluorescence and estimated K_d_(DFHBI-1T) and Fmax. (C) Correlations between measured fluorescence and estimated K_d_(Mg^2+^) and Fmax.

**Figure S11.**
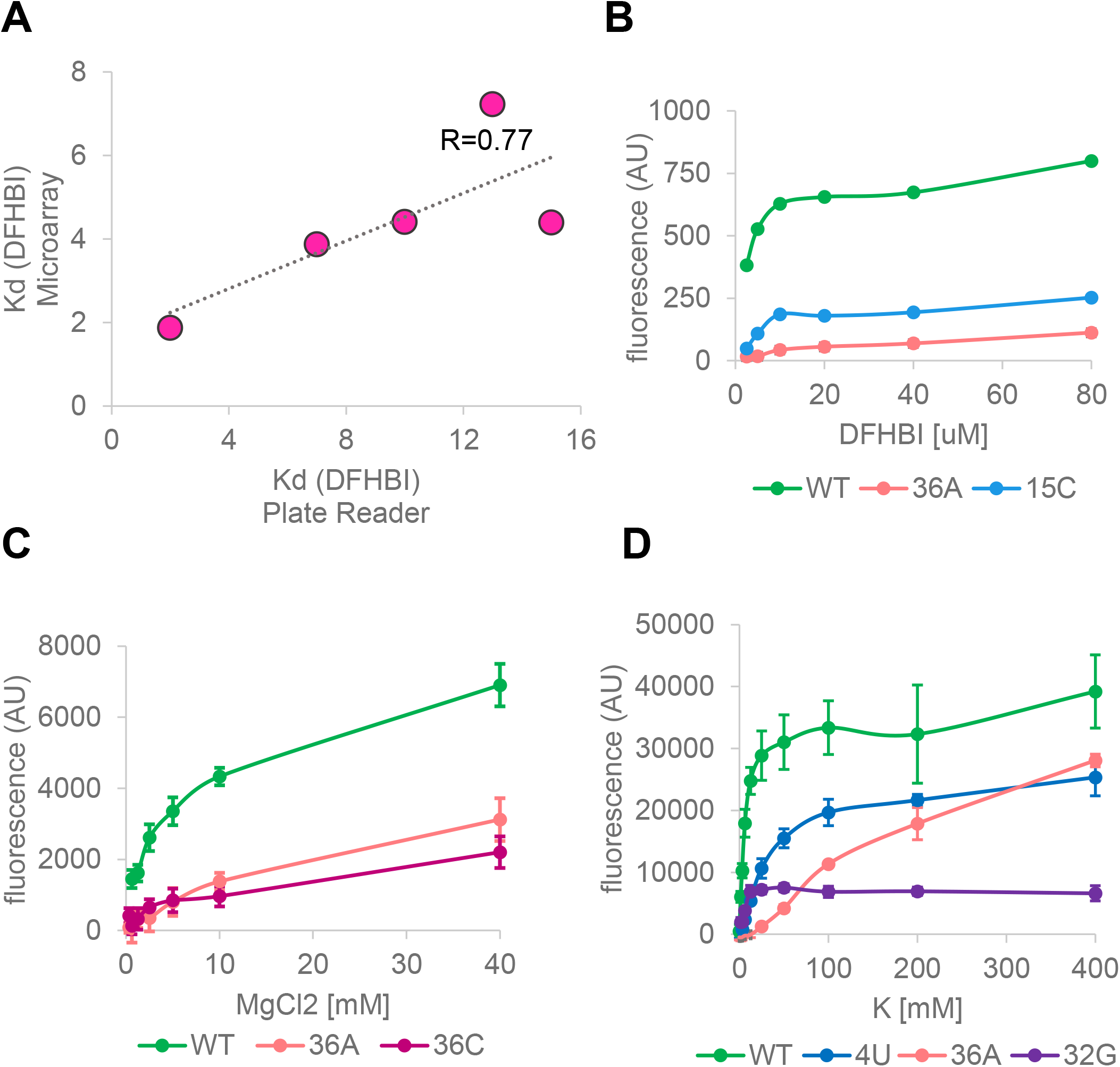
Spectrofluorometric analysis of dissociation rate constants. (A) Correlation of K_d_(DFHBI-1T) estimates from microarray and spectrofluorometric measurements for five selected variants. (B-D) Fluorescence of indicated Broccoli variants as a function of DFHBI-1T, Mg^2+^ and K^+^ concentration. Error bars represent standard deviation from 4 experimental replicates.

**Figure S12.**
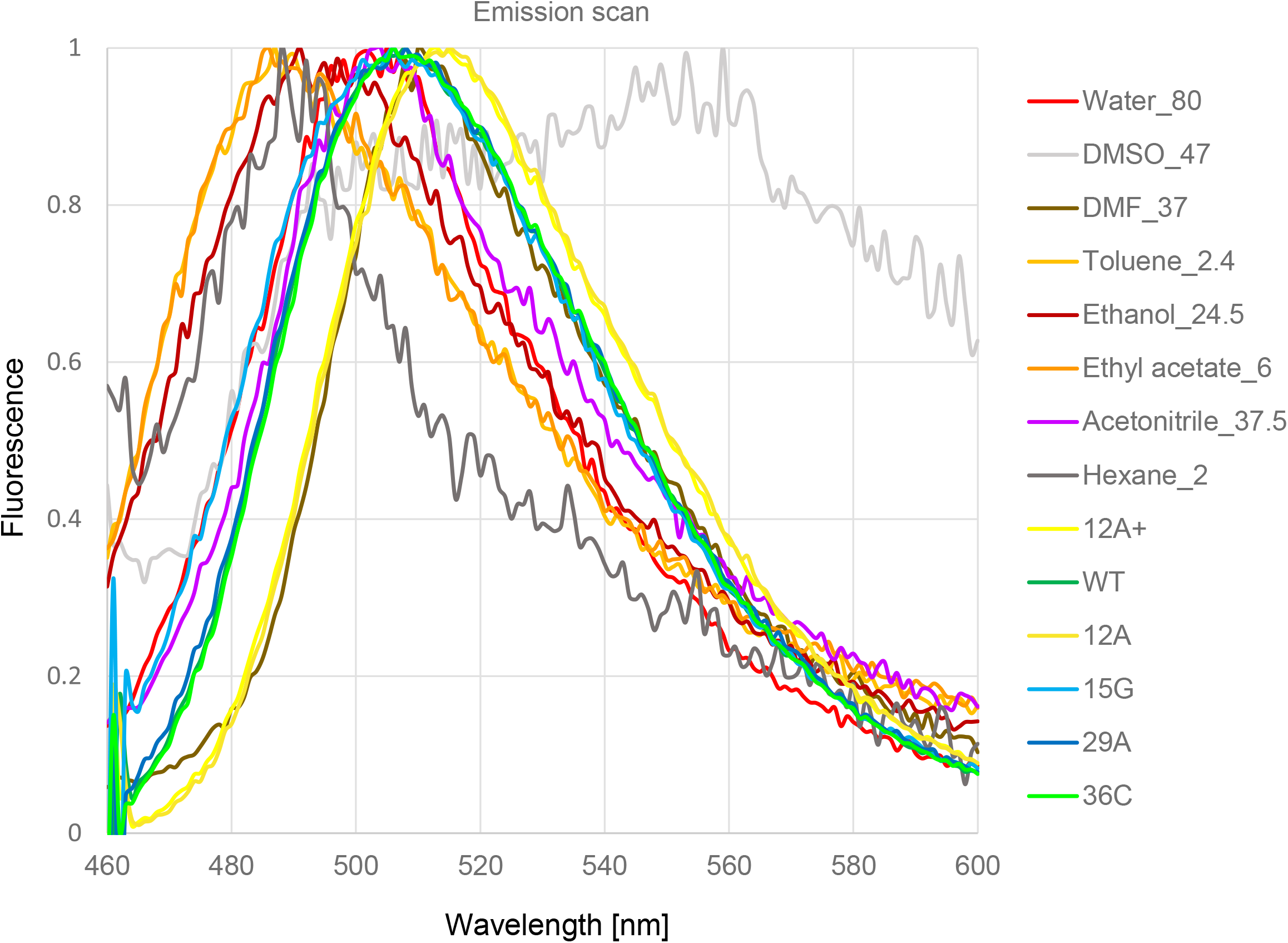
Emission spectra of DFHBI-1T in different solvents and when bound to selected variants of Broccoli in an aqueous buffer.

